# Cortical contributions to the perception of loudness and hyperacusis

**DOI:** 10.1101/2023.03.24.534013

**Authors:** Matthew McGill, Caroline Kremer, Kamryn Stecyk, Kameron Clayton, Desislava Skerleva, Kenneth Hancock, Sharon G. Kujawa, Daniel B. Polley

## Abstract

Sound perception is closely linked to the spatiotemporal patterning of neural activity in the auditory cortex (ACtx). Inhibitory interneurons sculpt the patterns of excitatory ACtx pyramidal neuron activity, and thus play a central role in sculpting the perception of sound. Reduced inhibition from parvalbumin-expressing (PV) inhibitory interneurons and the associated increased gain of sound-evoked pyramidal neuron spike rates are well-established consequences of aging and sensorineural hearing loss. Here, we reasoned that changes in PV-mediated inhibition would directly impact the perception of loudness. We hypothesized that ACtx PV activity could function as a perceptual volume knob, where reduced or elevated PV activity would increase or decrease the perceived loudness of sound, respectively. To test these hypotheses, we developed a two-alternative forced-choice loudness classification task for head-fixed mice and demonstrated that noise-induced sensorineural hearing loss directly caused a ∼10 dB loudness hyperacusis that begins hours after noise-induced sensorineural hearing loss and persists for at least several weeks. Conversely, sounds were perceived as ∼10 dB softer during optogenetic activation of ACtx PV neurons without having any effect on the overall detectability of sound. These data suggest that ACtx PV neurons can bi-directionally control the perceived loudness of sound, presumably via the strength of their inhibition onto local pyramidal neurons. Further, these data identify cortical PV neurons as a target for hyperacusis therapies and demonstrate a direct link between acquired sensorineural hearing loss and loudness hyperacusis.

## INTRODUCTION

Sensory systems encode the physical properties of environmental stimuli and reformat those features into spatially distributed neural activity patterns that form the basis of perception. The auditory system employs a variety of temporal, spatial, and population codes to represent the physical properties of sound (Chatterjee and Zwislocki, 1998; Joris et al., 2004; Micheyl et al., 2013; Pennington and David, 2020). The neural basis of loudness is derived from the physical intensity of sound and is directly supported by several proposed coding schemes, including the aggregate activity of a population of neurons, the selectivity of intensity-tuned neurons, and the location of responsive neurons (Schreiner and Malone, 2015; Zwicker and Fastl, 1999).

Loudness is the most fragile perceptual attribute of sound. In the case of the approximate 20% of adults with clinical hearing loss, low- and moderate-intensity sounds are perceived as inaudible or softer than normal, but the opposite side of the spectrum is also commonly observed; for an estimated 9-15% of adults with hyperacusis, moderate intensity sounds are perceived as disproportionately loud, unpleasant, or even painful (Andersson et al., 2002; Lin et al., 2011; Paulin et al., 2016). Loudness hyperacusis has been modeled in laboratory animals via noise-induced sensorineural hearing loss (Alkharabsheh et al., 2017; Hayes et al., 2014; Hickox and Liberman, 2014), and through systemic injections of highly concentrated salicylate, the active ingredient in aspirin (Auerbach et al., 2019; Zhang et al., 2014). Neural recordings from the central auditory pathway following any of these treatments or as a consequence of normal aging have identified a disproportionate increase in neural activity with increasing sound level, which has been coined “excess central gain” (Eggermont, 2017; Kotak, 2005; McGill et al., 2022; Noreña et al., 2003; Salvi et al., 2017). Excess central gain is most prominent at the level of the auditory cortex (ACtx) (Auerbach et al., 2014; Balaram et al., 2019; Chambers et al., 2016; Qiu et al., 2000). Sound-evoked hyper-responsivity in cortical pyramidal neurons arises in part from reduced inhibition, particularly from parvalbumin-expressing (PV) GABAergic interneurons that synapse onto the perisomatic compartment of pyramidal neurons to effectively throttle neural excitability and regulate action potential probability (Resnik and Polley, 2017; Sarro et al., 2008; Yang et al., 2011).

These observations suggest that the aggregate sound-evoked activity of ACtx excitatory neurons – particularly cortical projection neurons that innervate pre-frontal cortex, amygdala, striatum, and other targets outside of the central pathway - functions as a volume knob for loudness perception that can be effectively turned up and down by the strength of PV-mediated intracortical inhibition. A few key predictions of this model for loudness perception have been borne out by recent work. For example, recent studies have tracked the activity of ACtx neural ensembles before and after precisely controlled forms of hearing loss to show that PV activity levels (Resnik and Polley, 2021) and PV-mediated inhibition (Resnik and Polley, 2017) decline shortly after cochlear afferent input drops, just as hyper-responsivity, and hyper-synchrony of cortical excitatory neurons (McGill et al., 2022) emerges, particularly in long-range ACtx projection neurons (Asokan et al., 2018). The corollary hypothesis – that elevating the activity of ACtx PV neurons above normal would reduce the perception of loudness - has never been examined.

The largest missing element for an experimental proof of the neural basis of loudness perception and hyperacusis is that loudness itself has not been directly measured in laboratory animals with hearing loss but instead has only been studied with indirect proxies for loudness. Following noise exposure, acoustic startle responses are exacerbated (Hickox and Liberman, 2014) and operant Go-NoGo behaviors have shown increased growth slopes for sound level detection and sensitivity to direct stimulation of thalamocortical projection neurons (McGill et al., 2022). Likewise, reaction times in Go-NoGo tasks are reduced after salicylate administration (Auerbach et al., 2019). However, none of these are measures of loudness, *per se*, but instead assay behavioral reactivity to sound (Eggermont, 2013). On the other end of the spectrum, place preference behaviors have elegantly modeled the affective qualities of hyperacusis, such as auditory aversion or even pain, but they too provide no direct insight into loudness perception or loudness hyperacusis (Manohar et al., 2017).

To address this need, we developed a two-alternative forced-choice (2AFC) classification task in head-fixed mice that allowed robust, comparative measurements of loudness perception. We documented changes in loudness perception over several weeks after different methods of controlled cochlear injuries known to reliably induce ACtx hyperactivity. In separate cohorts of mice, we optogenetically activated ACtx PV neurons to demonstrate their direct influence on loudness perception.

## RESULTS

### Monotonic increase in reported loudness with increasing sound intensity

In humans, loudness perception and hyperacusis are evaluated using a variety of methods including a loudness contour test where subjects assign sounds to loudness categories (Cox et al., 1997; Jahn, 2022). We adapted the loudness categorization measurement in mice using a two-alternative forced-choice (2AFC) task, inspired by prior descriptions of frequency categorization in mice (Xin et al., 2019; Xiong et al., 2015). Mice were rewarded for licking the “soft spout” shortly following the presentation of low intensity 11.3 kHz tones (40-45 dB SPL) and for licking the “loud spout” shortly after the onset of higher intensity tones (75-80 dB SPL). Choice was measured over a 40 to 80 dB SPL range, where mice were conditionally rewarded for the low and high range but were rewarded upon selecting either spout for tones in the intermediate range (50-70 dB SPL; **Figure 1A**). This reward framework ensured that choice was strictly dependent on sound intensity without influencing how mice reported the transition between soft and loud sounds across an intermediate range of intensities.

**FIGURE 1.**
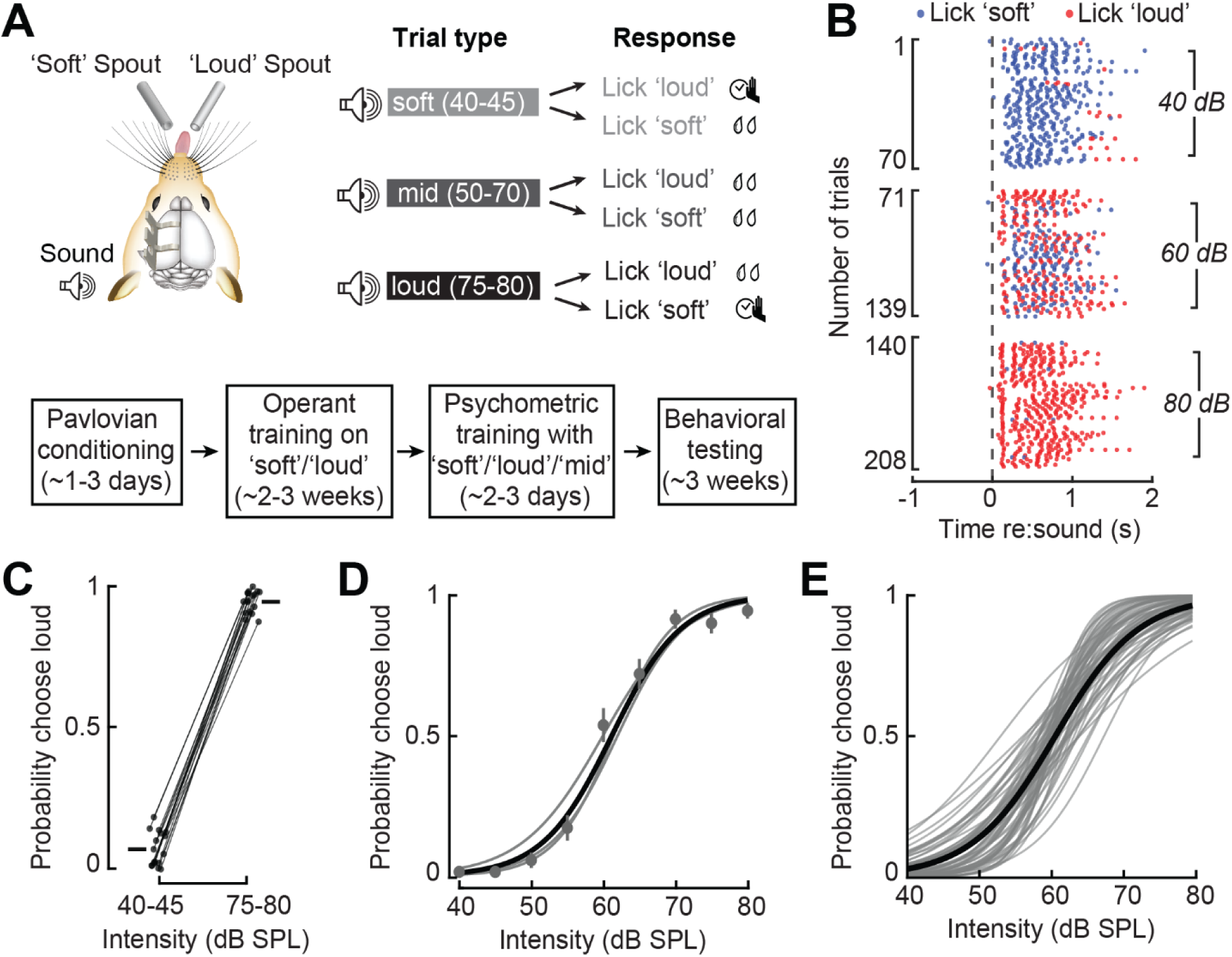
A two-alternative forced-choice task to assess loudness perception in mice. **(A)** *Top*, mice are trained to categorize 11.3 kHz tones as either ‘soft’ or ‘loud’. During testing, mice are required to correctly categorize 40-45 dB SPL (‘soft’) and 75-80 dB SPL (‘loud’) to receive reward but are allowed to choose either spout for mid-level intensities ranging from 50- 70 dB SPL. *Bottom*, an outline of the training timeline for this task. **(B)** Lick rasters from one example behavioral session for trials at three different intensities. Color represents spout choice. **(C)** The baseline choice probability at the extreme intensities for each mouse shows clear task-engagement with higher incidence of ‘loud’ choices for higher intensities compared to lower intensities (paired t-test, p = 2.8 x 10^-19^). **(D)** Baseline psychometric fit line. **(E)** Baseline behavioral choice curves from all mice (N=17 mice, 63 sessions), and the grand average.

Mice accurately reported sounds at the low and high ends of the continuum with short-latency licks on the soft and loud spout, respectively (**Figure 1B-C**). Mice alternately reported intermediate sound intensities (e.g., 60 dB SPL) as soft or loud from one trial to the next, yielding a smoothly varying transition in choice probability when averaged across trials (**Figure 1D**). Loudness reporting across behavioral sessions was highly reliable across all mice tested, with a 50% choice probability consistently occurring at approximately 60 dB SPL, the center of the intensity range (**Figure 1E**).

### Noise-induced cochlear sensorineural damage causes loudness hyperacusis

Noise exposure is among the most common risk factors for hyperacusis in humans (Fredriksson et al., 2022; Liberman et al., 2016; Muhr and Rosenhall, 2010). In animal models, exposure to intense noise has been linked to behavioral hypersensitivity to spared frequencies at the edge of the cochlear lesion (Hickox and Liberman, 2014; McGill et al., 2022) and increased neural gain in the inferior colliculus and ACtx (Chambers et al., 2016; McGill et al., 2022; Mulders and Robertson, 2013; Noreña et al., 2003; Qiu et al., 2000). To determine whether noise-induced hearing loss is associated with loudness hyperacusis at spared frequencies, mice were exposed to high-intensity noise (16-32 kHz at 103 dB SPL for 2 hr).

Auditory brainstem response (ABR) measurements performed one day and 14 days after noise exposure revealed a permanent threshold shift (PTS) for frequencies greater than 11.3 kHz (**Figure 2A**, left). To address the possibility that any behavioral changes after PTS could reflect pre-neural changes related to an increased spread of excitation along the basilar membrane or outer hair cell (OHC) damage, we exposed an additional cohort of mice to moderate intensity noise (8-16 kHz at 93 dB SPL for 2 hr) that caused a temporary threshold shift (TTS) one day after noise exposure that was completely reversed when tested again 14 days later (**Figure 2A**, right).

**FIGURE 2.**
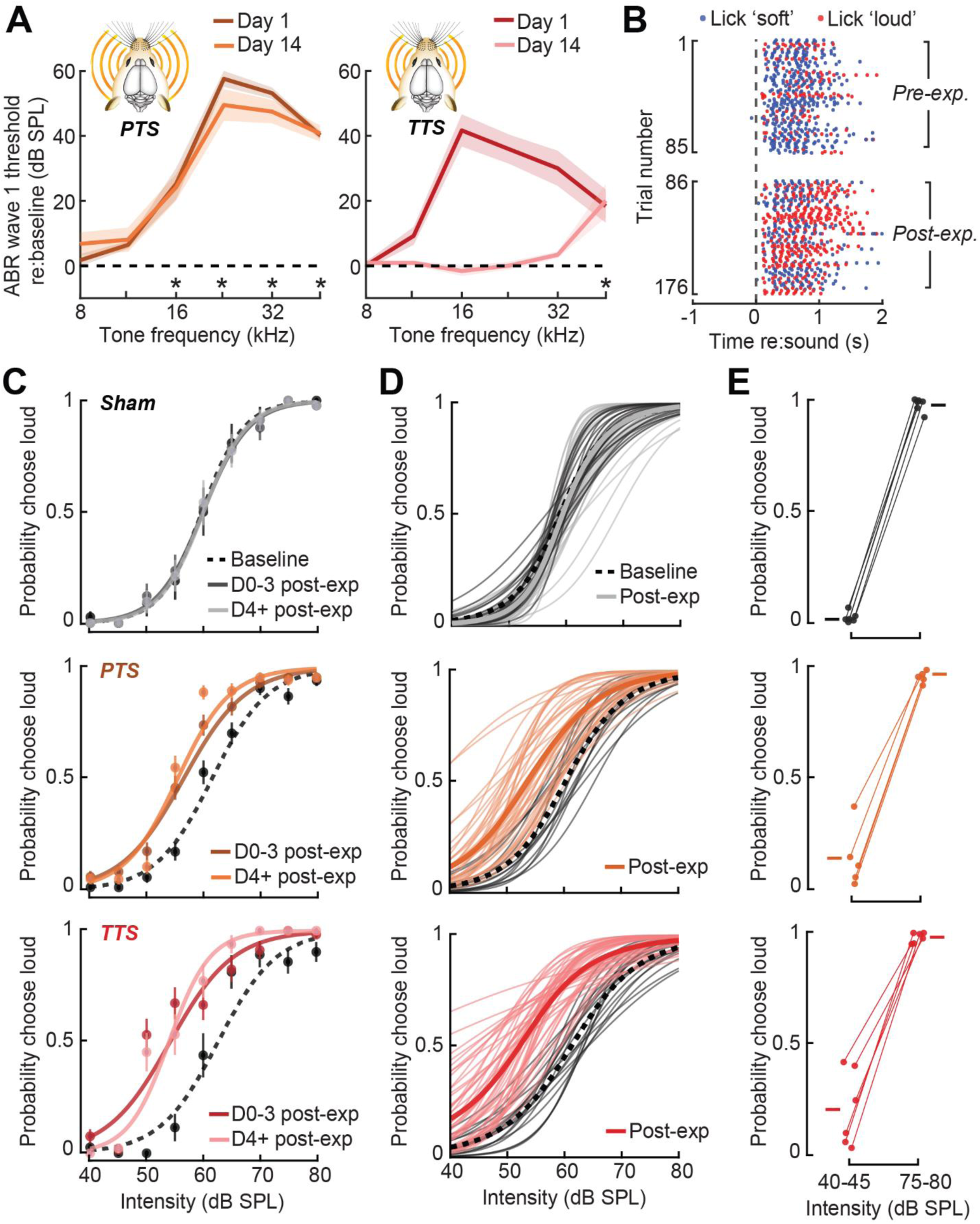
Following noise-induced hearing loss, mice show increased loudness reporting while maintaining proper task engagement. **(A)** Auditory brainstem response (ABR) wave 1 thresholds after PTS noise exposure shows a permanent threshold shift at high frequencies that’s stable at two weeks, while TTS noise exposure shows a general recovery of ABR thresholds after two weeks (2-way repeated measures analysis of variance [ANOVA] for PTS: Frequency term [F = 25.63, p = 7.6 x 10^-7^] and Time term [F = 0.33, p = 0.60], 2-way repeated measures ANOVA for TTS: Frequency term [F = 10.53, p = 1.6 x 10^-5^] and Time term [F = 47.2, p = 3.2 x 10^-10^]). Sham exposed mice did not show any change in ABR thresholds (data not shown). Asterisks denote significant differences between Day 14 and baseline ABR threshold with post hoc pairwise comparisons (p < 0.05). **(B)** Lick rasters from representative behavioral sessions of a noise-exposed mouse in response to a 55 dB SPL tone. The mouse shows increased ‘loud’ choices after noise exposure. **(C)** For one example mouse from each exposure group, the performance in baseline and across groups of days post-exposure. After both PTS and TTS, mice show increased loudness reporting across a range of intensities. Error on raw data points is bootstrapped standard error. **(D)** All baseline and post-exposure behavioral sessions (N=6/5/6, Sham/PTS/TTS, 187 total sessions), with average performance across mice before and after exposure shown in the darker lines. After both PTS and TTS, mice show a consistent leftward shift in their behavioral choice function, indicating increased loudness reporting. **(E)** The post-exposure choice probability at the extreme intensities for each mouse. After all exposure conditions, mice still display task engagement with higher incidence of ‘loud’ choices for higher intensities compared to lower intensities (paired t-test, Sham: p = 6.1 x 10^-8^, PTS: p = 1.4 x 10^-4^, TTS: p = 1.5 x 10^-4^).

Mice continued to produce appropriately timed lick behaviors after noise exposure, though intermediate tone intensities (e.g., 55 dB SPL) categorized more often as soft before exposure were subsequently reported as loud after exposure (**Figure 2B**). A relatively stable loudness hyperacusis following PTS and TTS was found both at the individual mouse level (**Figure 2C**) and was observed consistently across the performance of many mice and behavioral sessions (**Figure 2D**). Mice that underwent the same training and testing procedures simultaneously but were instead exposed to a non-damaging sound (40 dB SPL, 8-16 kHz for 2 hr), sham exposed mice, showed little change in task performance and demonstrated consistency in behavioral choice across time (Figure 2C-D, top row). Behavioral choices in noise-exposed mice remained faithful to task demands, in the sense that they continued to largely report low intensity sounds as soft and high intensity sounds as loud to receive conditional water reward (**Figure 2E**). The shift towards increased loudness was most apparent at intermediate sound levels, where choices were unconditionally reinforced, which is consistent with noise-induced hearing loss causing loudness hyperacusis.

To help better establish the link between induced cortical hyperactivity and behavioral hypersensitivity, we quantified changes in loudness choice across time and intensity. Following noise exposure, ACtx hyperexcitability and disinhibition have been shown to be stable across weeks (McGill et al., 2022; Resnik and Polley, 2021), setting up the hypothesis that loudness hyperacusis would be similarly stable over time. In support of this hypothesis, we found that TTS and PTS mice showed a stark increase in loudness choice on the first day after noise exposure that was stable across two weeks of behavioral testing and was more pronounced for lower mid-level intensities such as 55 dB SPL (**Figure 3A**). Effects at the highest intensities were likely smaller due to the already high probability of loud choices in baseline performance. The probability of reporting intermediate intensities as loud increased by 15.1% - 39.2% depending on sound intensity (**Figure 3B**). Interestingly, the percent increase in loud reporting was equivalently increased in the PTS and TTS cohorts and both were significantly greater than sham-exposed mice, suggesting that loudness hyperacusis is linked to sensorineural damage but not to the degree of threshold elevation.

**FIGURE 3.**
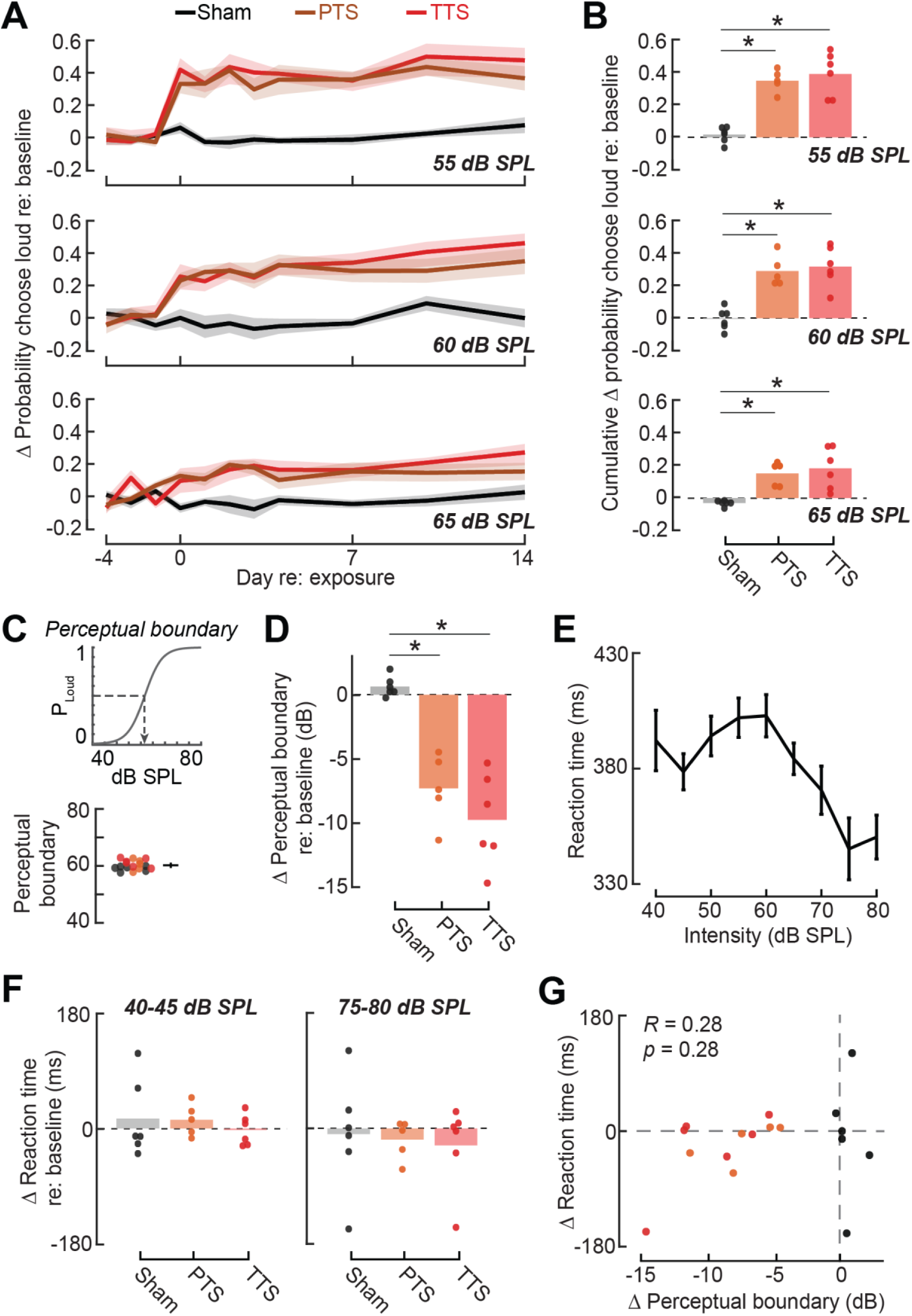
Loudness hyperacusis is stable across time but is not associated with decreased reaction time. **(A)** For each group, the mean change in the probability the mice chose ‘loud’ relative to their choice probability in baseline for three different mid-level intensities across time. Noise exposed mice show increased loudness reporting that is stable across time (mixed repeated-measures ANOVA with Group as a factor and Intensity and Time as repeated measures, Group x Intensity interaction term, F = 52.1, p = 3.3 x 10^-7^, and Time main effect, F = 2.3, p = 0.09). **(B)** For each intensity in (A), the change in choice probability per mouse, averaged across time. For each intensity, PTS and TTS groups show significantly increased loudness reporting after exposure compared to the sham group (t-test, p < 0.01, corrected for multiple comparisons), while the PTS and TTS groups show comparatively similar increases in loudness reporting (t- test, p > 0.5, corrected for multiple comparisons). **(C)** *Top*, we define the perceptual boundary as the intensity where the psychometric fit to the choice function crosses P_Loud_ = 0.5. *Bottom*, the boundary point in baseline for all mice. Each group showed a similar distribution of boundary points in baseline (1-way ANOVA, F = 3.0, p = 0.08). **(D)** The change in this perceptual boundary after exposure for each mouse. PTS and TTS groups show a similarly significant decrease in perceptual boundary compared to the sham group (1-way ANOVA, F = 25.8, p = 2.0 x 10^-5^, and for post-hoc pairwise comparisons, asterisks denote significant differences with p < 0.001). **(E)** The average trend in reaction time across all mice in baseline for each intensity. **(F)** The change in reaction time after exposure for each mouse at the extreme low and high intensities. Reaction times changes are highly varied and show no obvious trend across groups (mixed repeated-measures ANOVA with Group as a factor and Intensity as a repeated measure, Exposure x Intensity interaction term, F = 0.02, p = 0.98). Pairwise post-hoc comparisons show no significant differences between groups (p > 0.5). **(G)** For each mouse, the change in the perceptual boundary versus the change in reaction time at 75-80 dB SPL. There is no significant relationship between these variables (p = 0.28).

Increased loudness reporting after both PTS and TTS can also be quantified as a leftward shift in the psychometric choice function. To distill the loudness choice psychometric function down to a single value that is more easily compared with previous studies of auditory hypersensitivity, we defined the ‘perceptual boundary’ as the point where the fitted psychometric choice function crosses a probability of loud choice equal to 50% (**Figure 3C**). In baseline performance, the perceptual boundary was highly consistent across mice and groups with an average of 60.2 dB SPL (Figure 3C). The perceptual loudness boundary was effectively unchanged in sham-exposed mice compared with pre-exposure baseline but was significantly and equivalently shifted towards low sound intensities after PTS and TTS by an average of 8.6 dB, which amounts to more than a 7-fold change in signal amplitude (**Figure 3D**).

Previous studies have reported faster auditory reaction times in Go-NoGo operant tasks, which has been interpreted as evidence of hyperacusis (Auerbach et al., 2019; Zhang et al., 2014). In our 2AFC task, we noted that reaction times at baseline testing were approximately 50 ms faster for high-intensity sounds than low-intensity sounds, supporting the relationship between reaction time and perceptual salience (**Figure 3E**). However, we did not observe any significant or systematic changes in reaction time following PTS or TTS, even though these same mice showed evidence of loudness hyperacusis (**Figure 3F**). Moreover, there was no correlation across mice between the degree of shift in the loudness boundary and the change in reaction time (**Figure 3G**).

### There is no evidence for loudness hyperacusis in the degree and form of noise-induced cochlear sensorineural degeneration

To determine whether the degree or form of noise-induced changes in the early stages of auditory processing can account for loudness hyperacusis, we investigated cochlear histopathologic and ABR changes in PTS, TTS, and sham-exposed mice. The loss of synaptic contacts between inner hair cells and auditory nerve afferent fibers was assessed by quantitative immunolabeling of the pre-synaptic ribbon protein marker, CtBp2, in close apposition to the post-synaptic receptor marker, GluR2 (**Figure 4A**) (Kujawa and Liberman, 2009). As expected, PTS and TTS mice exhibited significantly reduced auditory nerve afferent innervation in the high-frequency base of the cochlea, though the degree of cochlear synaptopathy was greater in PTS than TTS ears. OHC loss was only observed in the extreme base of PTS ears (**Figure 4B**).

**FIGURE 4.**
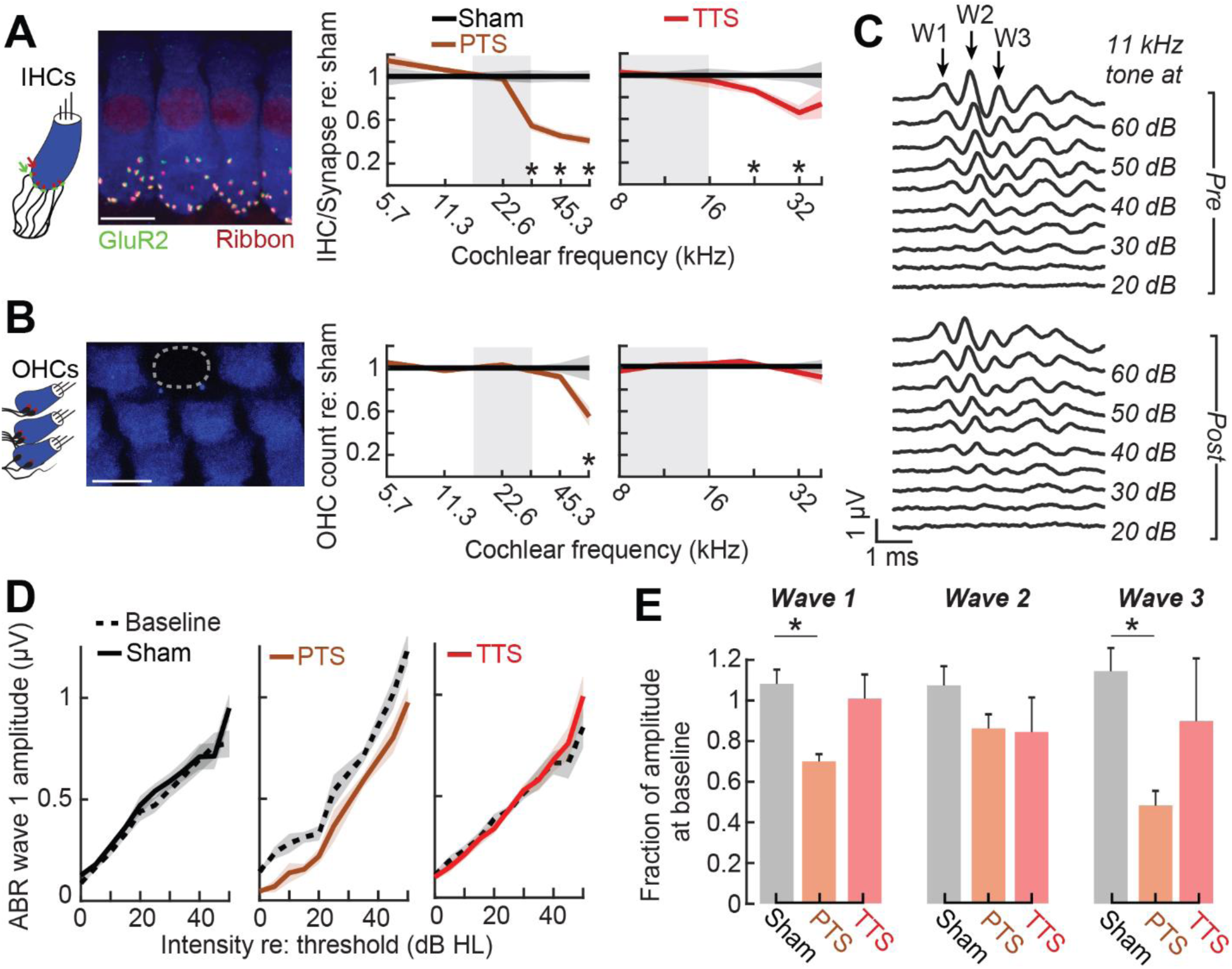
Measurements from the auditory periphery cannot account for loudness hyperacusis. **(A)** *Left*, cochlea immunostained for anti-CtBP2 and anti-GluR2a reveal presynaptic ribbon and post-synaptic glutamate receptor patches on inner hair cells (IHC’s). *Right*, both PTS and TTS exposures lead to a permanent reduction of IHC functional synapses in the high frequency region compared to sham-exposed cochleae processed simultaneously (mixed repeated measures ANOVA with Group as a factor and Frequency as a repeated measure for PTS: Frequency x Group interaction, F = 22.0, p = 4.3 x 10^-11^, and mixed repeated measures ANOVA for TTS: Frequency x Group interaction, F = 4.2, p = 0.003). Asterisks denote significant differences between sham and noise exposure with post hoc pairwise comparisons (p < 0.005). PTS data adapted from (McGill et al., 2022). Scale bars represent 10 µm. **(B)** *Left*, examination of OHC bodies. *Right*, loss of OHC’s is limited to the extreme basal regions of the cochlea only in the PTS group when compared to sham-exposed cochleae (mixed repeated measures ANOVA with Group as a factor and Frequency as a repeated measure for PTS: Frequency x Group interaction, F = 11.8, p = 2.8 x 10^-7^, and mixed repeated measures ANOVA for TTS: Frequency x Group interaction, F = 2.6, p = 0.09). Asterisks denote significant differences between sham and noise exposure with post hoc pairwise comparisons (p < 0.005). PTS data adapted from McGill, et al. (2022). Scale bars represent 10 µm. **(C)** Example pre- and post-exposure 11.3 kHz ABR waveforms at a range of intensities for one example noise-exposure mouse (W1 = Wave 1, etc.). **(D)** Average pre- and post-exposure ABR Wave 1 growth functions to 11.3 kHz tone pips across mice for each group. **(E)** Relative change in the ABR Wave 1-3 amplitude at 11.3 kHz, evaluated as the area under the ABR growth function compared to the same measure in baseline per mouse, for each exposure group. PTS and TTS groups do not show increased amplitude when compared to the sham exposure group, and in some cases mice that underwent PTS show hypoactivity. Asterisks denote significant pairwise differences (t-test, corrected for multiple comparisons, p < 0.01).

If loudness hyperacusis can be accounted for by excess gain at early stages of neural sound processing, one might expect that the amplitude of ABR waves would be elevated above normal levels. To test this hypothesis, we measured the amplitude of ABR waves elicited by the same 11.3 kHz pure tone frequency that was used in the loudness categorization task, for which there was no synaptopathy and minimal ABR threshold shift. The electrode montage we used for ABR recordings yielded waves 1, 2, and 3, which roughly correspond to the initial volley of synchronized action potentials of Type-I spiral ganglion neurons, cochlear nucleus neurons, and activity in the superior olivary complex and lateral lemniscus, respectively (**Figure 4C**) (Melcher et al., 1996).

Looking at Wave 1, ABR growth across mice is similar in sham and TTS exposed mice compared to baseline (**Figure 4D**). PTS exposed mice show a decrease ABR Wave 1 amplitude (Figure 4D), which is likely due to the slight threshold shift at 11.3 kHz (∼5 dB) observed in the ABR thresholds (Figure 2A). Expanding this analysis to all ABR Waves for 11.3 kHz, we computed the area under the ABR growth function after exposure relative to the same measure in baseline separately for each mouse and wave. TTS exposed mice showed no significant differences compared to sham exposed mice, while PTS exposed mice even showed a decrease in ABR amplitude compared to sham exposed mice for waves 1 and 3 (**Figure 4E**).

Thus, we found no evidence for hyper-responsivity in the early ABR waves in TTS or PTS mice at the same test frequency that was reliably associated with loudness hyperacusis in our 2AFC task. Although measurements from far-field recordings of gross electrical potentials under anesthesia do not rule out the possibility that excess central gain in early stages of neural processing can account for hyperacusis, the clear weight of evidence showing that central gain changes occur faster, last longer, and are expressed to an overall greater degree at later stages of processing supports the hypothesis that the most likely candidate for the neural underpinnings of loudness hyperacusis after sensorineural hearing loss would be found in the ACtx and, to a lesser extent, the inferior colliculus or medial geniculate body (Auerbach et al., 2014; Herrmann and Butler, 2021).

### Perceptual hyposensitivity during PV-mediated optogenetic silencing of ACtx

If the neural substrate for perceived loudness critically involved the ACtx, then ACtx should be necessary to perform the loudness categorization task. Cortical inactivation studies have demonstrated the necessary involvement of the ACtx in 2AFC categorization tasks that use more complex spectrotemporal stimuli (Guo et al., 2017; Znamenskiy and Zador, 2013), though ACtx is not required for Go-NoGo frequency recognition tasks that use pure tone stimuli (Ceballo et al., 2019). Here, the loudness categorization task uses pure tone stimuli but asks the animals to attend to the stimulus level, not frequency, and to indicate their percept with a 2AFC task design. As such, the loudness categorization task has some elements consistent with tasks that do require the involvement of ACtx but other features found in tasks that do not require the involvement of ACtx. A third possibility, which is the basis of the hypothesis tested here, is that suppressing the ACtx would not affect tone detection accuracy but would instead impact the accurate reporting of tone loudness.

The preferred method for inactivating the ACtx is to optogenetically activate GABAergic PV neurons as pyramidal neuron silencing can be interleaved with trials with normal pyramidal neuron signaling, thus affording a more direct comparison of behavior with and without cortical silencing than could be achieved with lesions or pharmacological inactivation (Atallah et al., 2012; Zhou et al., 2014). PV neurons are also hypothesized to be key regulators of loudness perception, given their functional role as throttles on the aggregate activity levels of ACtx pyramidal neurons. As described above, downregulation of PV activity and PV-mediated inhibition is an essential contributor to excess cortical gain and its perceptual sequelae of hyperacusis and tinnitus (Masri et al., 2021). The corollary hypothesis is that excess PV-mediated inhibition in the ACtx would cause sounds to be perceived as softer than would be expected based on their physical intensity. Based on this logic, the prediction stated above can be reformulated as silencing the ACtx via PV activation would not affect tone detection accuracy but would systematically shift loudness categorization towards reporting *hypo*acusis.

To test these hypotheses, we optogenetically activated PV GABAergic neurons in the left and right ACtx of *PV-Cre x Ai32* mice performing the 2AFC loudness categorization task. PV activation was performed on approximately 25% of trials that were randomly interleaved with laser off trials. PV activation was initiated 100 ms prior to tone onset and gradually tapered off following tone offset (**Figure 5A**). Mice remained task-engaged throughout behavioral sessions and showed proper loudness categorization on interleaved trials without laser stimulation (**Figure 5B**). As predicted, failures to report tone loudness (i.e., miss trials) were exceedingly rare, accounting for just 0.3% in laser off trials and 0.8% with ACtx silencing (**Figure 5C**). Therefore, in agreement with previous reports, the ACtx is not necessary for tone detection (Ceballo et al., 2019; Jenkins and Masterton, 1982). In addition to tone detectability, ACtx silencing, like noise exposure, did not cause any systematic change in reaction time at low or high intensities (**Figure 5D**).

**FIGURE 5.**
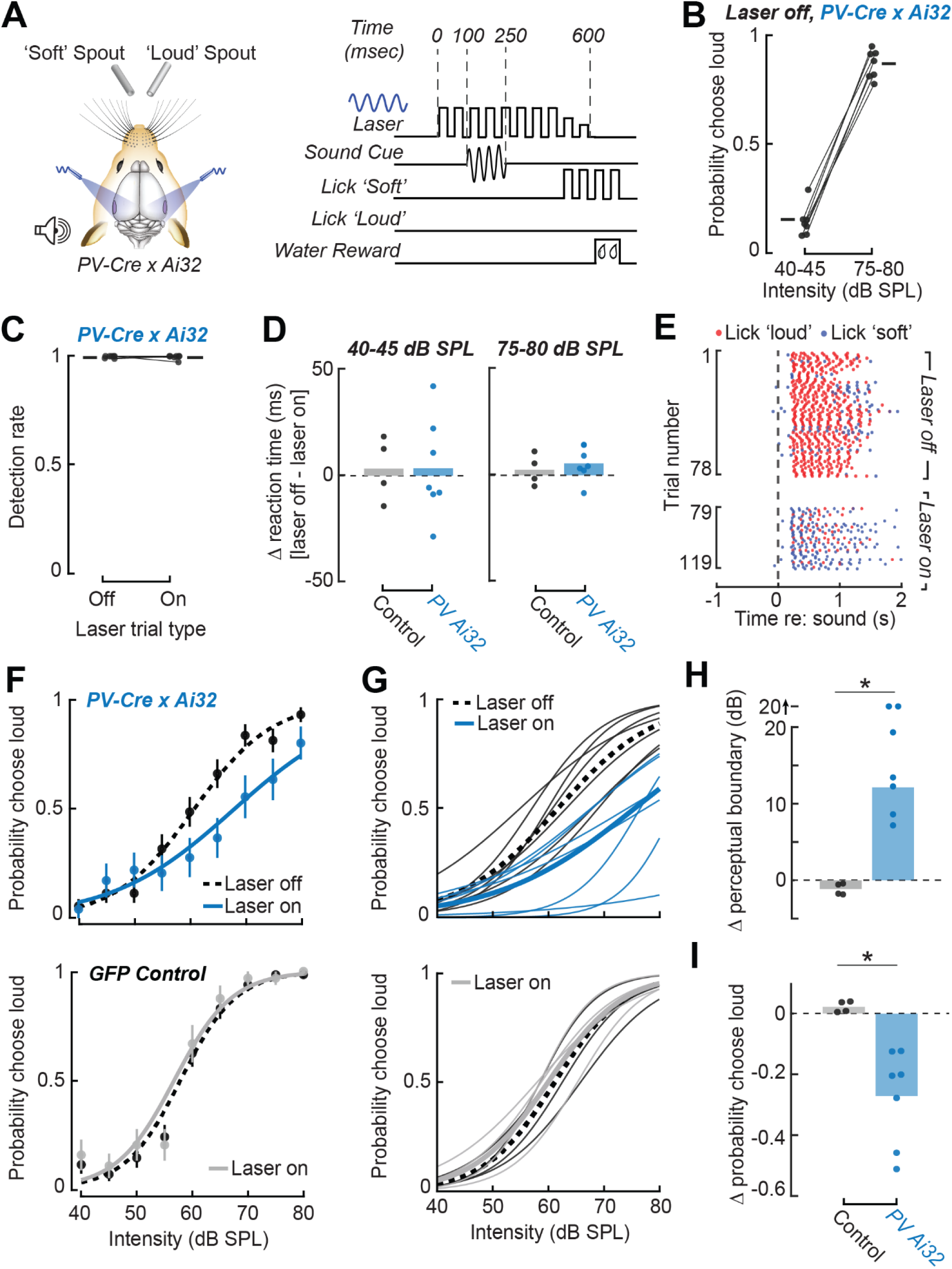
Inactivating ACtx via PV stimulation reduces perceived loudness. **(A)** *Left*, bilateral inactivation of ACtx using blue-light optogenetic activation of PV interneurons. *Right*, trial structure for interleaved trials of laser stimulation, which comprise ∼25% of trials. **(B)** Choice probability at the extreme intensities for each PV-Cre x Ai32 mouse without laser delivery. On trials without laser delivery, mice display task engagement with high incidence of ‘loud’ choices for higher intensities compared to lower intensities (paired t-test, p = 2.7 x 10^-7^). **(C)** The detection rate (the proportion of trials where a decision was provided) for all PV-Cre x Ai32 mice during both laser on and off trials. Mice do not show any difference in detection rate depending on trial type (paired t-test, p = 0.546). **(D)** The difference in the reaction time per mouse when the laser is on versus when the laser is off, separately evaluated at the extreme low and high intensities. Reaction times show no significant differences between mouse groups (t-test, 40/45 dB SPL: p = 0.50, 75/80 dB SPL: p = 0.27). **(E)** Lick rasters from a representative mouse in response to a 70 dB SPL tone with and without ACtx inactivation. The mouse shows increased ‘soft’ choices on trials with ACtx inactivation, but consistent reaction times. **(F)** Example behavioral choice functions with and without laser delivery for one PV-Cre x Ai32 mouse and one GFP-injected control mouse. The PV-Cre x Ai32 shows decreased loud reporting across a range of intensities while the GFP-injected control mouse shows consistent performance regardless of laser condition. **(G)** All mouse behavioral curves for each group (N = 7/3, Ai32/GFP) with average performance across mice shown in the darker lines both with and without laser delivery. GFP control mice show similar performance with and without laser delivery while PV-Cre x Ai32 mice show a rightward shift during PV-stimulation. **(H)** The change in the perceptual boundary per mouse when the laser is on versus when the laser is off. PV-Cre x Ai32 mice show a significant increase in their perceptual boundary compared to GFP controls, indicating decreased loudness choice (t-test, p = 4.7 x 10^-4^). For two mice, their psychometric line fit did not cross 50% and so their difference is denoted as ‘20^.’ **(I)** Since some mice did not have a measurable perceptual boundary, we instead quantified the mean difference in loudness choice probability across all intensities for each mouse. With this measure, PV-Cre x Ai32 mice show a significant decrease in their loudness choice probability compared to GFP controls (t-test, p = 0.0025).

Importantly, ACtx silencing via PV activation strongly biased the reported loudness of intermediate, unconditionally reinforced sound intensities towards soft. For example, a tone presented at 70 dB SPL was reliably classified as loud with the laser off but the percept switched to soft in interleaved trials from the same session with the laser on (**Figure 5E**). A shift towards soft reporting was observed across a wide range of higher intensities, both at the individual mouse level and across mice (**Figure 5F-G**, *top row*). As a negative control to show that hypoacusis was specific to PV activation and could not be accounted for by the addition of blue light or the expression of exogenous fluorescent proteins in PV neurons, we performed the same experiment on mice that expressed GFP in PV interneurons. However, optical excitation of GFP in PV neurons had no discernable effect on loudness reporting (Figure 5F-G, *bottom row*).

The overall rightward shift in the behavioral choice function with PV neurons activated amounted to an average 12.2 dB increase in the perceptual boundary, as compared to just an insignificant -1.1 dB change in GFP controls (**Figure 5H**). In some *PV-*Cre x *Ai32* mice, there was even no point at which the ‘laser on’ psychometric choice function crossed a 50% probability of choosing loud (Figure 5H). To address this issue, we also computed the average change in the probability of loudness choice across the intensity range when the laser is on compared to when the laser is off and observed that the probability of choosing the loud spout was reduced by an average of 27.2% during PV activation (**Figure 5I**).

### Modeling the contribution of task-dependent, stimulus-related, and experimental variables to mouse behavioral choice

Behavioral choices in a 2AFC task may be due to responses to the current stimulus, trial-history effects, and other behavioral strategies (Akrami et al., 2018; Busse et al., 2011; Lak et al., 2020; Mendonça et al., 2020). To pinpoint how the reporting of loudness (and changes therein after noise exposure or ACtx silencing) reflected the stimulus on that trial versus other co-variates, we created a logistic regression framework to predict single trial outcomes using a range of explanatory variables including the current trial intensity, the previous trial intensity, the previous trial choice, and all of their interactions (**Figure 6A**, see legend for variable coding).

**FIGURE 6.**
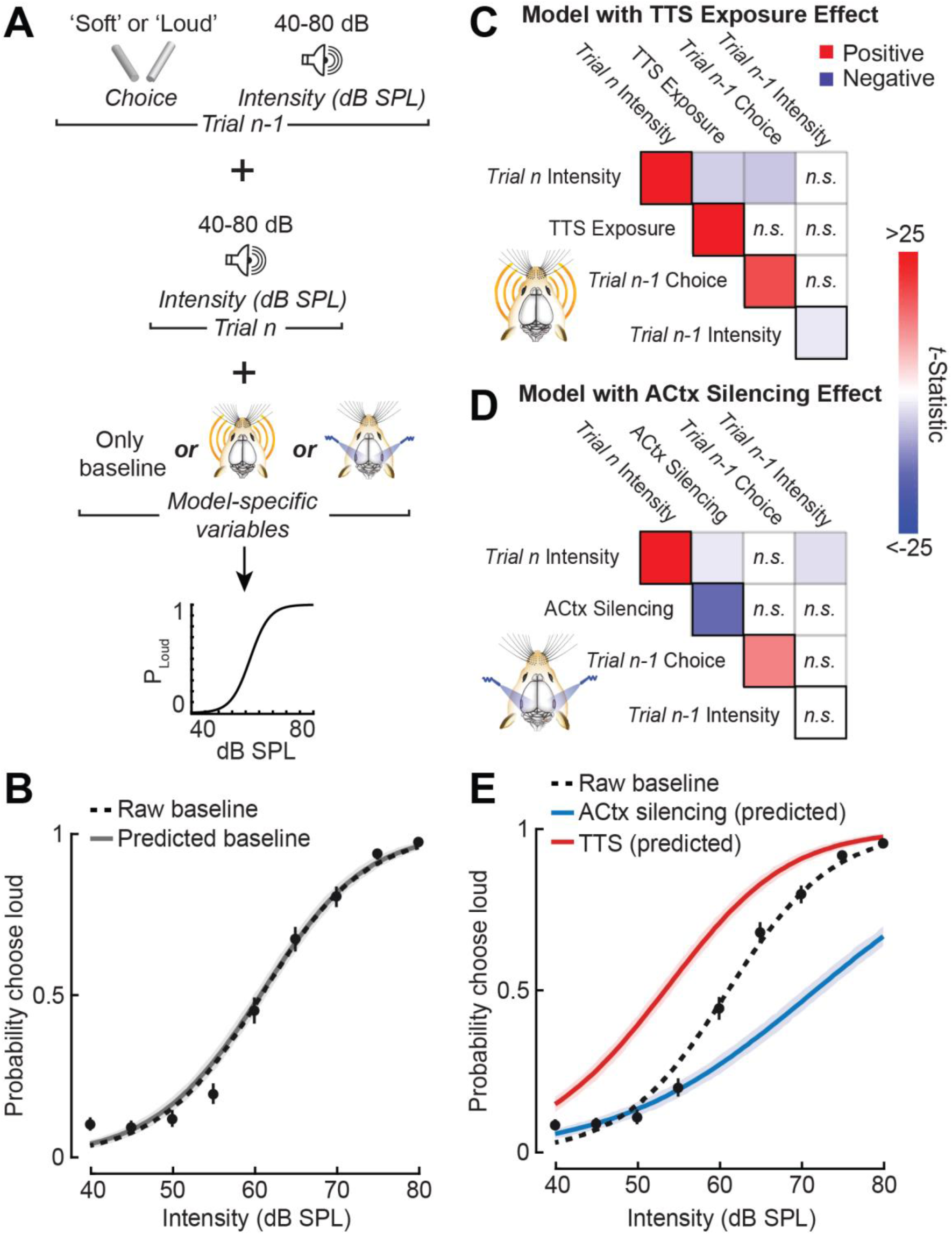
Noise exposure and ACtx inactivation induce opposing changes in loudness perception without affecting confounding mouse behavioral strategy. **(A)** Framework of a logistic regression model to predict mouse behavioral choice using: the left/right choice and intensity on the previous trial, the intensity on the current trial, and indications of experimental manipulations. Three separate models were constructed separately for the baseline performance, the noise exposure effect, and the optogenetic manipulation. All variables except intensity were coded as 0 vs. 1 indicator variables (i.e., 0/1 = choose soft/choose loud, 0/1 = sham/TTS exposure, 0/1 = laser on/laser off). **(B)** For the baseline model (Chi-square = 5.29 x 10^3^, p < 1 x 10^-299^), all raw baseline performance and the model-predicted performance with 95% confidence intervals. Raw and predicted functions show similar shape (model MSE = 0.12). **(C)** For the TTS exposure model (Chi-square = 1.38 x 10^3^, p = 1.2 x 10^-295^), the main effects for each variable (diagonal, black squares) and all pairwise interaction terms (off-diagonal). Color intensity indicates the magnitude of the t-statistic in the final model fit, and red/blue color indicates the relationship of each predictor variable with the response variable (i.e., increasing current trial intensity, TTS exposure, and previous trial choice of ‘loud’ all lead to increased [red] loudness reporting, while increasing the previous trial intensity leads to decreased [blue] loudness reporting). ‘n.s.’ indicates variables left out of the final model fit. TTS exposure does not interact with previous trial-related variables. **(D)** The same as (C) for the ACtx inactivation model (Chi-square = 1.13 x 10^4^, p < 1 x 10^-299^). Optogenetic silencing of ACtx does not interact with previous trial-related variables. **(E)** For the TTS exposure and the optogenetic silencing models, the predicted, isolated effects of TTS and ACtx silencing independent of all other behavioral variables. TTS exposure leads to increased loud reporting while ACtx silencing leads to increased soft reporting. Error represents 95% confidence intervals.

We constructed three separate behavioral models: one simply on normal baseline behavior, one for the post-exposure sessions in sham-exposed and TTS mice, and a third for behavior measured during the optogenetic experiment. Model-predicted baseline behavior was similar in psychometric function shape and overall loudness choice probabilities when compared to the raw mouse behavior (**Figure 6B**). Given our confidence in this framework to capture and replicate raw mouse behavior, we examined the fitted coefficients for each regression model examining either the effect of noise exposure or optogenetic silencing of ACtx.

Each regression model captured effects previously seen in the raw behavioral data and provided new behavioral insights. As expected, the current trial intensity accounted for the largest amount of behavioral choice variance and has a positive relationship with the probability of a loud choice (**Figure 6C-D**). TTS noise exposure and optogenetic silencing of ACtx both have large effects on loudness choice probability in opposing directions (Figure 6C-D).

Interestingly, mice show a procedural bias, reflected in the term for the previous trial choice, where a mouse is more likely to choose ‘loud’ after having previously chose the ‘loud’ spout and vice versa (Figure 6C-D). These logistic regression models also capture more modest effects, such as a trial-history effect where a louder sound intensity on the previous trial is correlated with a lower probability of choosing loud on the subsequent trial (Figure 6C). Most importantly, both TTS noise exposure and optogenetic silencing of ACtx do not interact with any trial-history variables (Figure 6C-D), indicating that our manipulations of ACtx excitability primarily affect perception of the current tone intensity rather than other latent behavioral variables or strategies (Figure 6C-D). Given these regression models, we can singularly extract the effect of either TTS noise exposure or ACtx optogenetic silencing by simulating behavioral trials where only these indicator variables are changed. The model-predicted effects of TTS and ACtx silencing highlights the bidirectional change in loudness perception compared to baseline behavioral performance where: (1) TTS noise exposure, which reduces PV activity, increases ACtx neural activity and causes loudness hyperacusis in the form of increased loud reporting, while (2) optogenetic elevation of PV activity, which reduces ACtx neural activity, causes loudness hypoacusis in the form of increased soft reporting (**Figure 6E**).

## DISCUSSION

Here, we introduced a 2AFC categorization task in head-fixed mice that allows for reliable assessment of loudness perception (Figure 1). In mice with synaptopathic or mixed sensorineural cochlear degeneration, sounds that were categorized as soft prior to noise exposure were subsequently reported as loud, demonstrating a causal link between cochlear afferent degeneration and loudness hyperacusis (Figure 2). A detailed characterization revealed that loudness hyperacusis: (i) occurs immediately following hearing loss, (ii) is stable across weeks, (iii) is most pronounced at mid-level intensities, and (iv) is not correlated with systematic changes in reaction time (Figure 3). Sensorineural degeneration was more extreme after noise that caused PTS compared to TTS, yet the degree of hyperacusis was equivalent between the two protocols with no observable antecedent of neural hyperactivity evident in the ABR (Figure 4). Categorizing the loudness of the tone – though not detecting the presence of the tone – required the ACtx and, more specifically, silencing ACtx via PV activation systematically biased mice towards reporting sounds as soft (Figure 5). A linear model found that manipulating central gain in opposite directions via noise-induced hearing loss and PV activation has a commensurate bi-directional effect on loudness perception without affecting other latent or task-related variables (Figure 6).

### Mechanistic insights

In ACtx, intensity-based information can be captured by the total number of active cells or their aggregate activity, the activation of particular intensity-tuned neurons, or the location of active cells within topographic space (Schreiner and Malone, 2015). Our findings suggest that gross manipulations of ACtx excitability have a bi-directional effect on the perceived loudness of a delivered sound, which most directly supports an aggregate population code for the mapping of neural activity onto a loudness percept. Previous studies in frequency categorization have shown that targeted manipulations of ACtx population activity can bias the information received by downstream areas from ACtx-projection neurons and subsequently affect behavior (Guo et al., 2017; Xiong et al., 2015; Znamenskiy and Zador, 2013). Hyperexcitability in ACtx following noise-induced hearing loss has been shown to affect downstream areas through ACtx projections (Asokan et al., 2018), and so we would similarly expect these changes to bias behavioral choice as has been shown in our 2AFC task.

Implicit in our findings is the idea that the strength of PV-mediated inhibition may play a role in the perceived loudness of a sound. After hearing loss, there is decreased feedforward PV-mediated inhibition onto pyramidal neurons (Resnik and Polley, 2017; Yang et al., 2011), downregulation of GABA receptors (Balaram et al., 2019; Caspary et al., 2013; Sarro et al., 2008), and decreased PV neuron responses to sound in ACtx (Resnik and Polley, 2021). This disinhibition may drive loudness hyperacusis, and PV neuron disruption has already been implicated in other hearing related disorders such as tinnitus (Masri et al., 2021). Our previous work has found that pyramidal neurons with properties resembling weaker inhibitory strength are most susceptible to hyperactivity following noise exposure, underscoring the importance of inhibitory network strength in ACtx plasticity and resulting auditory hypersensitivity (McGill et al., 2022). Similar phenomena have also been reported in the somatosensory system, where both spared nerve injury and autism spectrum disorder have been associated with cortical disinhibition and an accompanying hypersensitivity to touch (Cichon et al., 2017; Klintwall et al., 2011; Robertson and Baron-Cohen, 2017). Furthermore, manipulations of PV and SOM neuron activity have also been implicated in other auditory tasks such as the detection of sound from background noise (Christensen et al., 2019; Lakunina et al., 2022), a task that is also disrupted by disinhibition induced through sensorineural damage in the cochlea (Resnik and Polley, 2021). Altogether, PV neurons may play a direct role in how ACtx population responses reflect sound intensity and how this activity is relayed to downstream areas to influence a loudness percept.

### Behavioral implications and limitations

The demonstration of a clear loudness hypersensitivity phenotype in laboratory animals is crucial for studies of the behavioral consequences and neurological underpinnings of sensorineural hearing loss or other related neurological conditions. In the auditory field, there is a frequent reliance on the use of reaction times in operant detection tasks or other indirect measures as proxies of loudness hyperacusis (Auerbach et al., 2019; Rybalko et al., 2015; Sun et al., 2012; Zhang et al., 2014). Here, we observe evidence of loudness hyperacusis based on mouse decision making in our 2AFC task, but we do not see faster reaction times or a correlation between the changes in reaction time and loudness choice. Consequently, reaction time may be limited as an indicator of hyperacusis and may be dependent upon the method of inducing hearing loss, the paradigms and timescales used to test the animals, or even the choice of animal model (Auerbach et al., 2019; Zhang et al., 2014). It’s important to note that a similar 2AFC loudness categorization task has been assessed in rats following noise-induced hearing loss (Alkharabsheh et al., 2017). However, in this study noise exposures were carried out before animals were trained to categorize the loudness of a tone, and so it is difficult to disentangle the contributions of various development- and hearing loss-related plasticity to perception. Our behavioral paradigm effectively allows for: (i) direct assessment of loudness perception, (ii) longitudinal measurements where single animals can serve as their own reference point, and (iii) pairing with optogenetic and other useful methodologies.

Hyperacusis encompasses both an increased loudness perception and affective components such as sound aversion and even pain (Tyler et al., 2014). Our 2AFC task is designed to measure the loudness component without providing any insight into the pain or distress components of hyperacusis. There are recent studies making progress assessing aversive components of sound following hearing loss using freely moving behavior (Manohar et al., 2017), and, importantly, these affective components of sound perception can affect its perceived loudness (Florentine et al., 2011). Although we are not directly measuring the effects of stress or other emotional components on sound perception, we controlled for changes in stress and pain related to all experimental procedures other than sensorineural hearing loss through the inclusion of a sham exposed group that underwent the same surgical, behavioral, and experimental procedures (Figures 2-3). Through our behavioral modelling, we did not find any interactions between hearing loss and other task-related variables, supporting the idea that behavioral changes solely reflected changes in loudness categorization (Figure 6). Sound hypersensitivity may be broader than the single frequency we chose to test, 11.3 kHz. Our methods of inducing cochlear synaptopathy are frequency-specific (Figure 4), and so the resulting ACtx hyperactivity is topographic (Noreña et al., 2003; Yang et al., 2011), where hyperactivity is most pronounced at frequencies near the edge of the cochlear lesion (McGill et al., 2022; Rajan, 1998). So, the choice of 11.3 kHz allows for both robust and thorough longitudinal assessment of loudness perception across a wide intensity range.

### Therapeutic implications

The cortical representation of sound intensity is plastic. ACtx responses to sound intensity can be modulated through learning and reward (Polley, 2006; Polley et al., 2004), or through neuromodulation such as the pairing of sound with cholinergic stimulation (Froemke et al., 2013). This plasticity is reflected behaviorally such as through a heightened ability to detect softer sounds or to discriminate sound intensities (Froemke et al., 2013; Polley, 2006; Polley et al., 2004), or to enhance cortical encoding of low SNR sounds in background noise (Whitton et al., 2014). These findings raise the possibility of harnessing endogenous brain plasticity mechanisms to suppress sound responsiveness and reduce loudness perception in subjects with auditory hypersensitivity. Here, our gross optogenetic inhibition of ACtx on single trials underscores the effectiveness of population activity manipulations on loudness perception (Figure 5). If PV hypofunction is an underlying cause of loudness hyperacusis, then reinvigorating PV-mediated inhibition onto cortical pyramidal neurons would provide a plausible means of remediating loudness hyperacusis. This might be best accomplished with gene therapy approaches to enhance cortical inhibitory tone (Masri et al., 2023) or with comparatively “low-tech” behavioral approaches that leverage the endogenous plasticity mechanisms of the adult brain to reduce the psychoaffective burden of auditory perceptual disorders if not directly reduce the salience of the unwanted percept itself (Polley and Schiller, 2022).

## MATERIALS AND METHODS

### Model and subject details

All procedures were approved by the Massachusetts Eye and Ear Animal Care and Use Committee and followed the guidelines established by the National Institute of Health for the care and use of laboratory animals. Data were collected from 41 mice. A total of 28 mice contributed to behavioral tasks: 17 to 2AFC post noise-exposure studies (N = 6/5/6, sham/PTS/TT) and 11 to optogenetic 2AFC behavior (N = 4/7, control/ChR2). A total of 13 mice contributed to in-depth auditory brainstem response (ABR) testing and cochlear histology (N = 7/6, sham/TTS). Optogenetic silencing and optogenetic control experiments were performed in PV Cre x Ai32 and PV Cre mice (JAX stock no: 017320 and 024109 respectively). All other behavioral experiments were performed in C57BL/6J mice (JAX stock no: 000664). Mice of both sexes were used for this study. Noise exposure occurred at 10 weeks postnatal and was timed to occur in the morning, when the temporary component of the threshold shift is less variable. Mice were maintained on a reverse 12 hr light/12 hr dark cycle and were provided with *ad libitum* access to food and water unless they were undergoing behavioral testing, in which case they had restricted access to water in the home cage.

### Survival surgeries for awake, head-fixed experiments

Mice were anesthetized with isoflourane in oxygen (5% induction, 1.5%–2% maintenance). A homeothermic blanket system was used to maintain body temperature at 36.6° C (FHC). Lidocaine hydrochloride was administered subcutaneously to numb the scalp. The dorsal surface of the scalp was retracted and the periosteum was removed. The skull surface was prepped with etchant (C&B metabond) and 70% ethanol. For behavioral subjects not tested with optogenetic manipulations, a titanium head plate (iMaterialise) was then affixed to the dorsal surface with dental cement (C&B metabond).

For mice undergoing additional optogenetic testing, the skull overlying the ACtx of each hemisphere was made optically transparent. To do so, one layer each of first super glue (Krazy Glue), then clear dental cement (C&B metabond), and nail polish (L.A. colors) were each applied and allowed to dry. While the nail polish was drying, small black plastic casings were placed over ACtx (Doric). For optogenetic control mice, GFP was expressed in PV neurons by making two small burr holes in the skull overlying the ACtx of each hemisphere (1.75-2.25 mm rostral to the lambdoid suture) with a 31-gauge needle. Pulled glass micropipettes (Wiretrol II, Drummond) were backfilled with virus solution, AAV2/5-hSyn-EGFP (addgene no. 27056), and 200 nL were injected into the targeted brain areas at 1.1 nL/s 0.3 mm below the pial surface using a precision injection system (Nanoject III, Drummond) with a 5 s delay between each injected bolus. At least 10 minutes passed following each injection before the pipette was withdrawn and burr holes were filled with KWIK-SIL (WPI). Injections were performed prior to the skull clearing procedure.

Once all elements had dried, a custom titanium headplate was affixed to the dorsal surface with dental cement, as per the procedure described above. At the conclusion of the headplate attachment for all surgeries, Buprenex (0.5 mg/kg) and meloxicam (0.1 mg/kg) were administered, and the animal was transferred to a warmed recovery chamber.

### High-frequency noise exposure

For TTS exposure, octave-band noise at 8-16 kHz was presented at 93 dB SPL for 2 hours. For PTS exposure, octave-band noise at 16-32 kHz was presented at 103 dB SPL for 2 hours. Exposure stimuli was delivered via a tweeter (Fostex USA) fixated inside a custom-made exposure chamber (51 x 51 x 51 cm). The walls of the acoustic enclosure were slanted such that no two walls were parallel as to minimize standing waves. Additionally, irregular surface depths and sharp edges were built onto 3 of the 6 walls using stackable ABS plastic blocks (LEGO) to diffuse the high-frequency sound field. Prior to exposure, mice were placed, unrestrained, in an independent wire-mesh chamber (15 x 15 x 10 cm). This chamber was placed at the center of a continuously rotating plate, ensuring mice were exposed to a more uniform sound field. Sham exposed mice underwent the same procedure except that the exposure noise, 8-16 kHz octave-band noise, was presented at an innocuous level (40 dB SPL). All sham and noise exposures were performed at the same time of day.

### Cochlear function tests

Animals were anesthetized with ketamine (120 mg/kg) and xylazine (12 mg/kg), with half the initial ketamine dose given as a booster when required, and they were placed on a homeothermic heating blanket during testing. ABR stimuli were 5 ms tone pips at 8,12,16 or 32 kHz with a 0.5 ms rise-fall time delivered at 27 Hz. Intensity was incremented in 5 dB steps, from 20–100 dB SPL. ABR threshold was defined as the lowest stimulus level at which a repeatable waveform could be identified. DPOAEs were measured in the ear canal using primary tones with a frequency ratio of 1.2, with the level of the f2 primary set to be 10 dB less than f1 level, incremented together in 5 dB steps. The 2f1-f2 DPOAE amplitude and surrounding noise floor were extracted. DPOAE threshold was defined as the lowest of at least two continuous f2 levels, for which the DPOAE amplitude was at least two standard deviations greater than the noise floor. All treated animals underwent rounds of DPOAE and ABR testing one week before noise- or sham-exposure, and immediately following the conclusion of behavioral or imaging testing.

### Cochlear histology

Deeply anesthetized animals were transcardially perfused with 4% paraformaldehyde in 0.1 M phosphate buffer, the cochleae were dissected and perfused through the round and oval windows and post-fixed in the same solution for 2 h at room temperature. Cochleae were then decalcified in 0.12 M EDTA for 2 days at room temperature, dissected into half-turns and blocked in 5% normal horse serum and 0.3% Triton X-100 for 1 hour at room temperature. They were immunostained by overnight incubation with a combination of the following primary antibodies: 1) rabbit anti-myosin VIIa (Proteus Biosciences, 1:200); 2) mouse IgG1 anti-CtBP2 (BD Biosciences, 1:200); 3) mouse IgG2a anti-GluA2 (Millipore, 1:2 000) and secondary antibodies AlexaFluor 647 donkey anti-rabbit, 1:200; Alexa Fluor 568 goat anti-mouse IgG1, 1:1 000; Alexa Fluor 488 goat anti-mouse IgG2a, 1:1 000 (Thermo Fisher Scientific) coupled to the blue, red, and green channels, respectively. Immunostained cochlear pieces were measured, and a cochlear frequency map was computed (Müller et al., 2005) to associate structures to relevant frequency regions using a plug-in to ImageJ (Parthasarathy and Kujawa, 2018).

Confocal z-stacks at 8.4, 16, 22.6, 32 and 45 kHz frequency locations were collected using a Leica TCS SP8 microscope using a glycerol-immersion objective (63x, N.A. 1.3) and 2.38x digital zoom to visualize inner hair cells (IHCs) and synaptic structures. Two adjacent stacks were obtained at each target frequency spanning the cuticular plate to the synaptic pole of ∼10 hair cells (in 0.33 µm z-steps). Images were loaded into an image-processing software platform (Amira; ThermoFisher Scientific), where IHCs were quantified based on their Myosin VIIa-stained cell bodies and CtBP2- stained nuclei. Presynaptic ribbons and postsynaptic glutamate receptor patches were counted using 3D representations of each confocal z-stack. Juxtaposed ribbons and receptor puncta constitute a synapse, and these synaptic associations were determined by calculating and displaying the x–y projection of the voxel space within 1 µm of each ribbon’s center (Liberman et al., 2011). OHCs were counted based on the myosin VIIa staining of their cell bodies.

### Two-alternative forced-choice task

Three days after headplate surgery, animals were weighed and placed on a water restriction schedule (1 mL per day). During behavioral training, animals were weighed daily to ensure they remained above 80% of their initial weight and examined for signs of dehydration. Mice were given supplemental water if they received less than 1 mL during a training session or appeared excessively dehydrated (e.g., fur tenting). During testing, mice were head-fixed in a dimly lit, single-walled sound-attenuating booth, with their bodies resting in an electrically conductive cradle. Tongue contact on the lickspout closed an electrical circuit that was digitized (at 40 Hz) and encoded to calculate lick timing. Target tones were delivered through an inverted dome tweeter (Parts Express) positioned 10 cm from the left ear. Digital and analog signals controlling sound delivery and water reward were controlled by a PXI system with custom software programmed in LabVIEW (National Instruments). Free-field sound levels were calibrated with a wide-band ultrasonic acoustic sensor (Knowles Acoustics).

Mice required approximately 3 weeks of behavioral shaping before they could perform the complete 2AFC task. For shaping, mice were first habituated to head-fixation and conditioned with a Pavolvian approach in which 11.3 kHz tones (150ms, 5 ms raised cosine onset-offset ramps) at 40 and 45 dB SPL were presented with a small quantity (3.5 µL) of water on the “soft” spout while 75 and 80 dB SPL tones were paired with water on the “loud” spout.

The assignment of loud and soft to the left and right spouts were randomly determined for each mouse. Trials had a variable inter-trial interval (4-10s) selected on each trial from a truncated exponential distribution to produce a flat hazard function. Tone presentation was subject to an additional potential delay until a 1s criterion for no licking was met. Once mice were rapidly licking the correct spout shortly following each tone they were converted to an operant task, in which water delivery was contingent upon licking the correct spout within 1.2s following tone onset. To forestall or mitigate the development of choice bias, a closed-loop training paradigm was intermittently used where mice were repeatedly given the same trial type after an incorrect response until they chose correctly.

Once mice met the criterion of > 90% correct categorization, they were advanced to the full stimulus set, where tone intensity on each trial was randomly selected from a 40 to 80 dB SPL (5 dB step size). At this stage, mice were required to correctly categorize 40-45 dB SPL tones as ‘soft’ and 75-80 dB SPL tones as ‘loud’ by licking the appropriate spouts to receive water, but they were rewarded regardless of their spout choice for 50-70 dB SPL tones. Mice were acclimated to the psychometric form of the task for 2-3 days, and then behavioral testing began.

All mice underwent the same training procedure. Mice used for noise- and sham-exposure testing underwent 3-4 days of baseline data collection and were then tested every 1-2 days following exposure for 2 weeks. Mice used for both bilateral optogenetic inhibition of ACtx and optogenetic controls were tested for 4-5 days following the completion of their training in order to collect an appropriate number of trials.

### Optogenetic stimulation

Trials with sound and light stimulation (25%) were randomly interleaved with sound-only trials. Blue light was delivered to the brain via an optic fiber/ferrule assembly (0.2 mm diameter, 0.22 NA Doric) coupled to a diode laser (Omnicron LuxX 473nm, 10mW at fiber tip, 25 ms pulse width at 20 Hz over a 600 ms period with a linear-tapering amplitude envelope for the last 100 ms to reduce any rebound excitation in ACtx), and the light path was split to provide a light source for each hemisphere separately. A black mating sleeve (Doric) was attached to each laser patch cable, which fit snugly into the skull-mounted plastic casings. Laser onset preceded tone onset by 100 ms.

### Behavioral analysis and modeling

Psychometric functions were fit to raw left/right choice data using binary logistic regression. To get the fits across time or across mice, raw data was first concatenated. The perceptual boundary was defined as the intensity where the psychometric function crossed a 50% loudness choice probability.

To model mouse behavioral choice, a generalized linear model was used with a log link function. Observations for model fitting were individual trials, where the response variable was the mouse’s choice on a trial coded as 0 = soft choice and 1 = loud choice. The predictor variables shown were the mouse’s choice on the previous trial (1 = loud choice), the intensity on the previous trial, and the intensity on the current trial. Three separate models were fit, one to the baseline data across all mice, one to noise exposure data which included one extra predictor indicator variable (1 = TTS), and one to all optogenetic inhibition data which included one extra predictor indicator variable (1 = laser on). Final model fits were found via a stepwise approach where a full model including all pairwise interactions terms was made, and then the least significant terms were removed from the model one-by-one until all predictor terms were significant. Model fits were made using ‘fitglm’ in MATLAB.

### Statistical analysis

All statistical analyses were performed in MATLAB 2017b (Mathworks). Details of all statistical testing are provided in the figure legends. Post hoc pairwise comparisons were corrected for multiple comparisons using the Holm-Bonferroni correction. Unless otherwise stated, error shown is the standard error of the mean (SEM).

## Author Contributions

MM and DP designed the research with input from KC and SK. MM, CK, and KS performed the behavioral experiments, MM performed the electrophysiological experiments, and DS performed the cochlear histology with guidance from SK. KH programmed the software for the behavioral training and data collection. MM analyzed all data, and MM and DP prepared the figures. MM and DP wrote the manuscript with input from all authors.

## Acknowledgements

These studies were supported by NIH grant DC009836 (DP), DC015857 (DP and SK), and NIH fellowship DC018974-02 (MM).

We thank Y. Watanabe for their guidance with surgical preparations for optogenetic experiments, and E. Smith and C. Rutagenwa for their assistance with cochlear histology.

